# Reconciling author names in taxonomic and publication databases

**DOI:** 10.1101/870170

**Authors:** Roderic D. M. Page

## Abstract

Taxonomic names remain fundamental to linking biodiversity data, but information on these names resides in separate silos. Despite often making their contents available in RDF, records in these taxonomic databases are rarely linked to identifiers in external databases, such as DOIs for publications, or ORCIDs for people. This paper explores how author names in publication databases such as CrossRef and ORCID can be reconciled with author names in a taxonomic database using existing vocabularies and SPARQL queries.

## 1 Linking taxonomic names

We can represent “core” biodiversity data as a network of connected entities, such as taxa and their names, publications, people, species, macromolecular sequences, images, and natural history collections [9]. Creating a “biodiversity knowledge graph” is an implicit goal of several initiatives in biodiversity informatics. Indeed, taxonomic databases were early adopters of the Resource Description Framework (RDF) for describing entities and their interrelationships. From 2005 onwards, major databases of taxonomic names (“nomenclators”) for plants, animals, and fungi have used Life Science Identifiers (LSIDs) [2] to uniquely identify those names. LSIDs can be dereferenced to return metadata in RDF [8], and several databases used the same vocabulary (developed by TDWG) to encode information about taxonomic names, their status (e.g., were the names in current use), and where the names were published. The use of globally unique identifiers that can be dereferenced, and which return data in a consistent, machine-readable format would seem to satisfy the preconditions for creating biodiversity knowledge graph [9].

Despite the obvious desirability of linking biodiversity data together ([1]), the biodiversity knowledge graph as yet to spontaneously assemble itself. Arguably the biggest reason is that there were few, if any, connections between taxonomic information and external data sources. For example, taxonomic databases typically cite the taxonomic literature using text strings, rather than persistent identifiers. Hence, we still have silos, albeit silos available in linked data formats.

Breaking those silos requires making links between taxonomic names and other entities of interest, such as the taxonomic literature, specimens, and taxonomists themselves. Creating these links is currently labour intensive, hence early efforts at constructing knowledge graphs have either had modest taxonomic and geographic scope (Ozymandias, [11]), or are closely linked to the output of one science publisher (OpenBioDiv, [12]).

In this paper I discuss some ways to combine data from taxonomic and publication databases to help seed a biodiversity knowledge graph. In this paper the focus is on plant names, but the ideas apply to names for other taxonomic groups, such as animals and fungi.

## 2 Populating the knowledge graph

### 2.1 Plant names

The Index of Plant Names Index (IPNI^1^) is an international register of published plant names based at the Royal Botanic Gardens, Kew but which has contributions from the Harvard Gray Index and the Australian Plant Name Index [3]. Both new taxonomic names (e.g., for newly described species) and new combinations (e.g., reflecting transfers of species from one genus to another) are recorded in IPNI, together with a citation to the scientific work that published that name. Increasingly these names are being submitted to IPNI during the publication process for a paper, rather than simply being captured after the fact [13]. Each name in the IPNI database has a LSID that uniquely identifies that name. I used the IPNI API to retrieve the corresponding RDF (in XML), fixed a small bug in the XML, then convert it to *n*-triples and uploaded it to a triple store (Blazegraph^2^).

### 2.2 Publications

IPNI contains terse citations for the publication of each name in its database. For some records the IPNI curators have added a link to the corresponding page in the Biodiversity Heritage Library (BHL^3^), and for some recently added records the IPNI web site may give the DOI for a publication, but the majority of IPNI records are not linked to a digital identifier for the publication associated with each name.

As part of ongoing work to match citation strings for taxonomic names to persistent identifiers for the corresponding publications ([10]), I developed a set of scripts to matching the text string citations to digital identifiers such as DOIs, Handles, JSTOR links, etc. The difficult part of this work is mapping the page-level citations stored in IPNI to work-level bibliographic data. Given a complete bibliography of the taxonomic literature, this would be a relatively straightforward task, in that we could treat each work as comprising a set of pages, and we simply ask which works include the page in the IPNI citation. However, as yet we don’t have a comprehensive bibliography of life ([6]), hence much of the work in making the mapping involves scouring the web for sources of bibliographic information in the hope that these will include works containing the IPNI citations. This mapping between IPNI names is periodically uploaded to a GitHub repository^4^, and has also been published as a ”datasette” [10]. For this project I took this mapping and exported it in RDF.

Metadata about the publications themselves publications was harvested from CrossRef^5^ and ORCID ^6^, expressed as RDF using terms from the Schema.org^7^ vocabulary, and added to the knowledge graph.

### 2.3 People

I used the same approach to modelling authors employed in Ozymandias [11], namely using the schema:Role type [5]. Inserting a Role node between two entities enables us to annotate that relationship, for example specifying the time period during which the relationship applies. Rather than directly connect a publication to its creator, the creator of a work is a Role, which in turn has the author as its creator property. We can then store the position of author in the list of authors using the schema:roleName property (e.g., “1”, “2”, etc.).

Although CrossRef does include some author ORCIDs, in many cases it does not. We can match CrossRef and ORCID records using DOIs for publications, and augment CrossRef records with ORCIDs (see Fig. 1). Using a SPARQL query (see Listing 1.1) and a DOI we can match the same author names and order of authorship (role) from CrossRef with those for the same record in ORCID. This query assumes that the author names are identical in the CrossRef and ORCID, an obvious refinement would be to accept approximate matches.

**Fig. 1.**
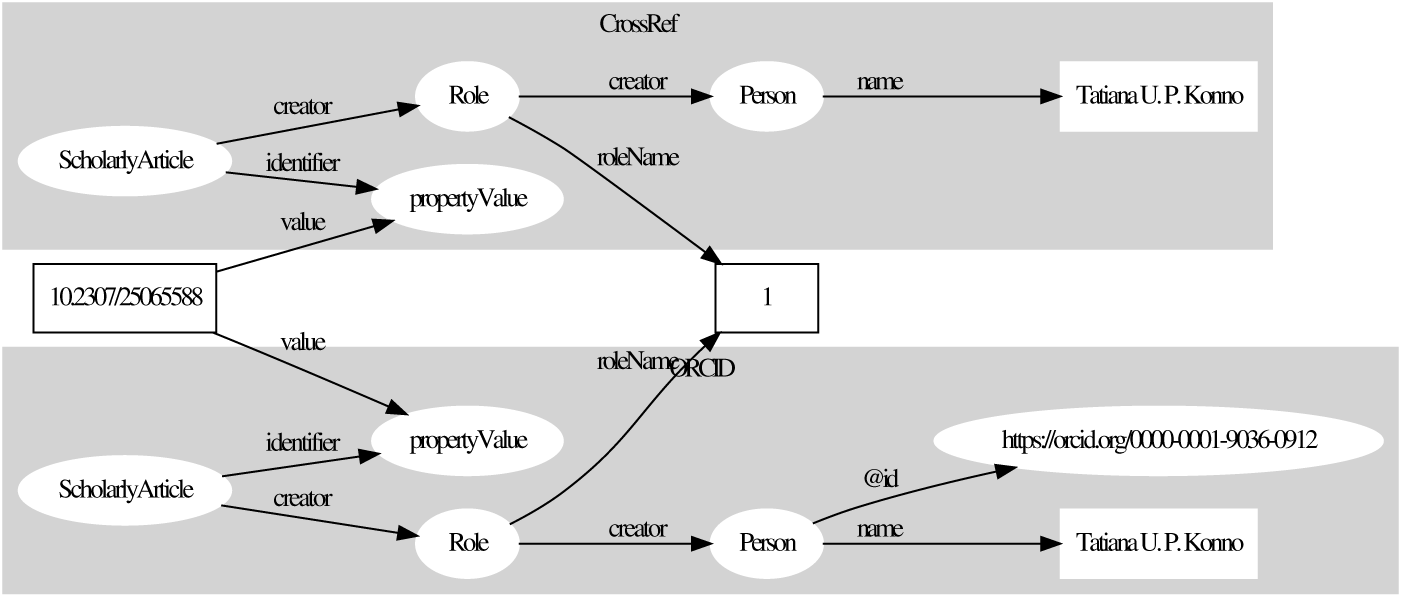
Matching metadata from CrossRef and ORCID for the article with the DOI 10.2307/25065588. The author “Tatiana U. P. Konno” in CrossRef has the ORCID identifier “0000-0001-9036-0912”.

IPNI has a similar, if more complex notion of “roles”. For each taxonomic name there is a “team” of one or more authors, each of whom may have various roles, such as author of the original name, or author of a subsequent version (“combination”) of the name (for example, the change in name that results when a species moves from one genus to another). IPNI records the role of each team member (e.g., whether they are a “publishing author” or a “combination author”) and their position in an ordered list of team members. Hence one approach to matching publication authors (and associated identifiers) to IPNI authors (and their IPNI identifiers) is by matching on roles (Fig. 2). Given this model we can find candidates for matching IPNI team members to publication authors using a SPARQL query (Listing 1.2).

**Fig. 2.**
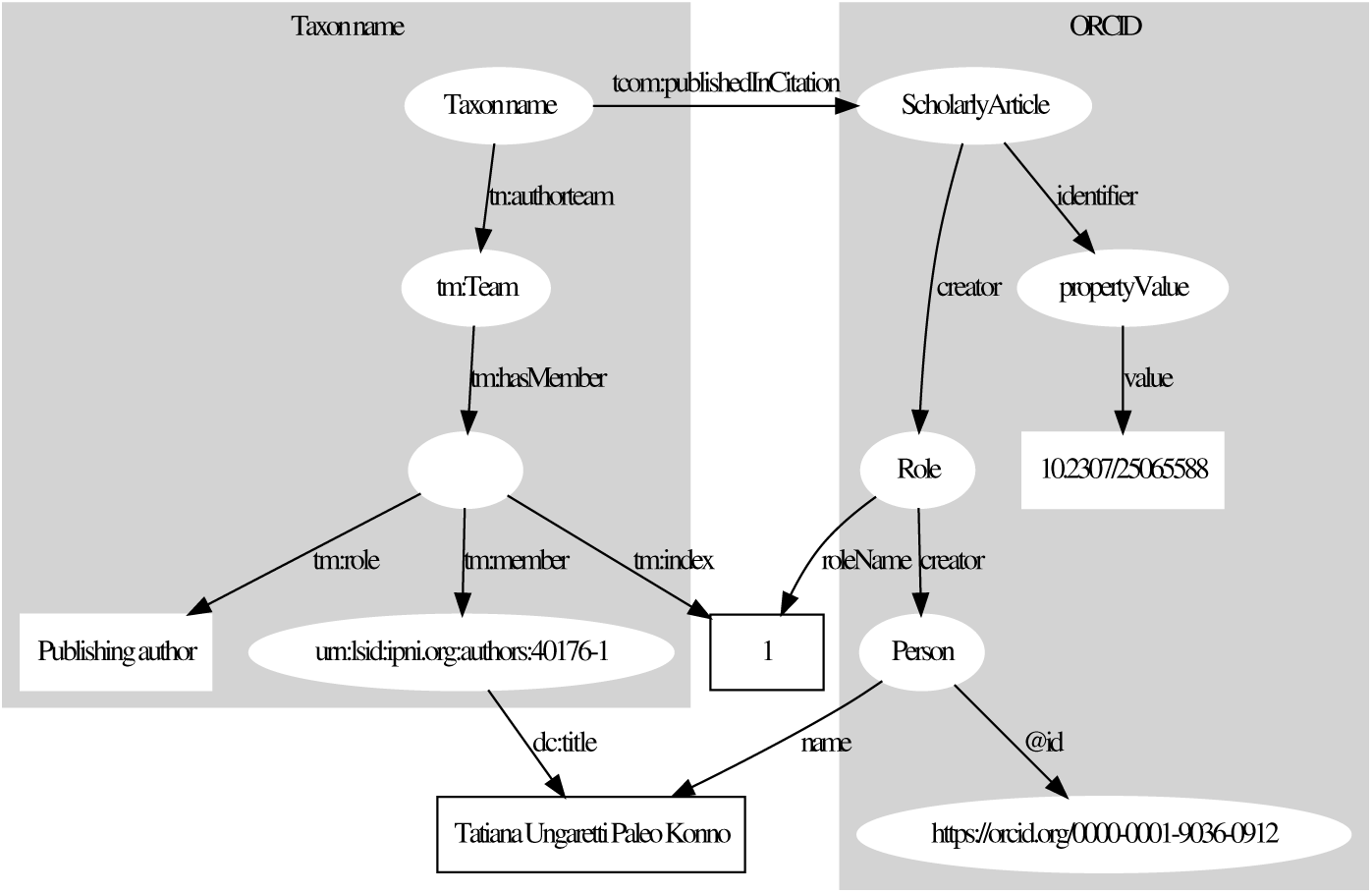
Matching a member of the author team for a plant name to the author of the paper publishing that name.

This query finds matches between the first author of the paper with DOI 10.2307 / 25065588 [7] and the first member of the team that coined a plant name *Minaria cordata* (Turcz.) T.U.P.Konno & Rapini (LSID urn:lsid:ipni.org:names:77074588-1). The author is Tatiana Ungaretti Paleo Konno (ORCID 0000-0001-9036-0912).

### 2.4 Problems and limitations

In the process of assembling the plant name knowledge graph I have encountered several limitations of the methods outlined here. For example, the order of authorship author in a taxonomic team and in a publication need not be the same. It is not uncommon for an ORCID profile to list only the owner of that profile as the author of a publication, and IPNI only records authors who actually name the plant, which may not be all the authors of corresponding publication. Hence we cannot completely rely on the orderings to make the mapping.

Relatively few taxonomists have ORCID profiles, and these profiles are often incomplete. Indeed of the 483 ORCID profiles currently in the knowledge graph, only 367 include publications, leaving over 100 profiles that cannot be matched to the taxonomic literature using the type of query shown in Fig. 2. The queries described here rely on exact string matching and hence will fail if the source databases do not have the same name for the same person. Currently 6965 IPNI taxonomists in the knowledge graph are linked to a publication with full bibliographic metadata, but only 2101 of those can be matched to one of the authors of that publication. Improving on that matching will require approximate string comparison.

The proposed mapping of authors to identifiers could be also used to augment existing database records, such as Wikidata^8^ [15]. At the time of writing (24 Sep 2019), there is a Wikidata item Q16300981 (revision 880704436) for Tatiana Ungaretti Paleo Konno. This item has IPNI author id 40176-1 as an attribute, but not an ORCID id. Alessandro Rapini (a coauthor on [7]) has two Wikidata entries (Q54703172 and Q5574227). The current revision of Q54703172 (revision 1009574566) lists the IPNI author ID for Alessandro Rapini, but not the ORCID, whereas Q54703172 (revision 974791158) has an ORCID but not an IPNI id. Hence we have two entries for the same author. Wikidata is an obvious venue for storing author identifiers, however many of these identifiers are added to Wikidata using automated “bots” which speed up data entry but don’t necessarily ensure that the entity they are adding is not already in Wikidata.

## 3 Summary and future directions

The approach outlined here is merely a first step in fleshing out a knowledge graph of plant names. By connecting names to publications we provide more details on the provenance of those names. Taxonomic publications, especially recent ones, are likely to have details on aspects of the morphology, ecology, and even genomics of the species being considered. By converting text string citations into identifiers such as DOIs, we also open up the possibility of linking the paper to citing literature, or to additional data from that publication, such as DNA sequences stored in GenBank^9^, and evolutionary trees stored in TreeBASE^10^.

There are extensive programs to digitise natural history collections, and the results of digitisation are being aggregated into global databases such as the Global Biodiversity Information Facility (GBIF)^11^. However, data aggregators have been criticised for not giving attribution and credit to individual researchers such as taxonomists [4]. By associating author names with identifiers such as ORCIDs we can link taxonomists additional outputs of their research, such as new species descriptions or new taxonomic classifications. Generating lists of taxonomists and associated identifiers can also facilitate linking those researchers to specimens in natural history collections that they have collected or identified, further enhancing our ability to document the activities of taxonomists [14].

## 4 Listings

**Listing 1.1.**
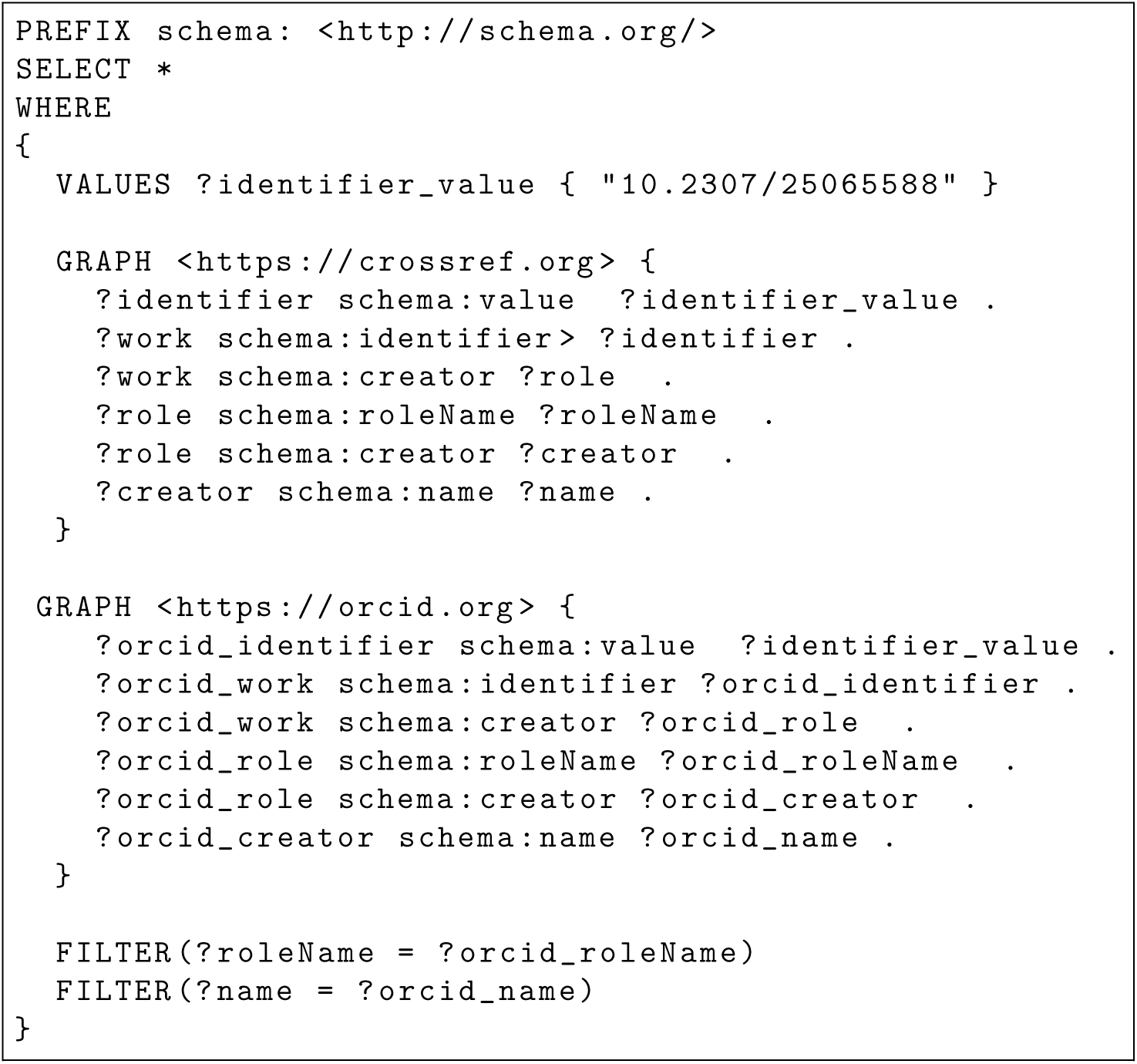
SPARQL query to match authors in CrossRef and ORCID

**Listing 1.2.**
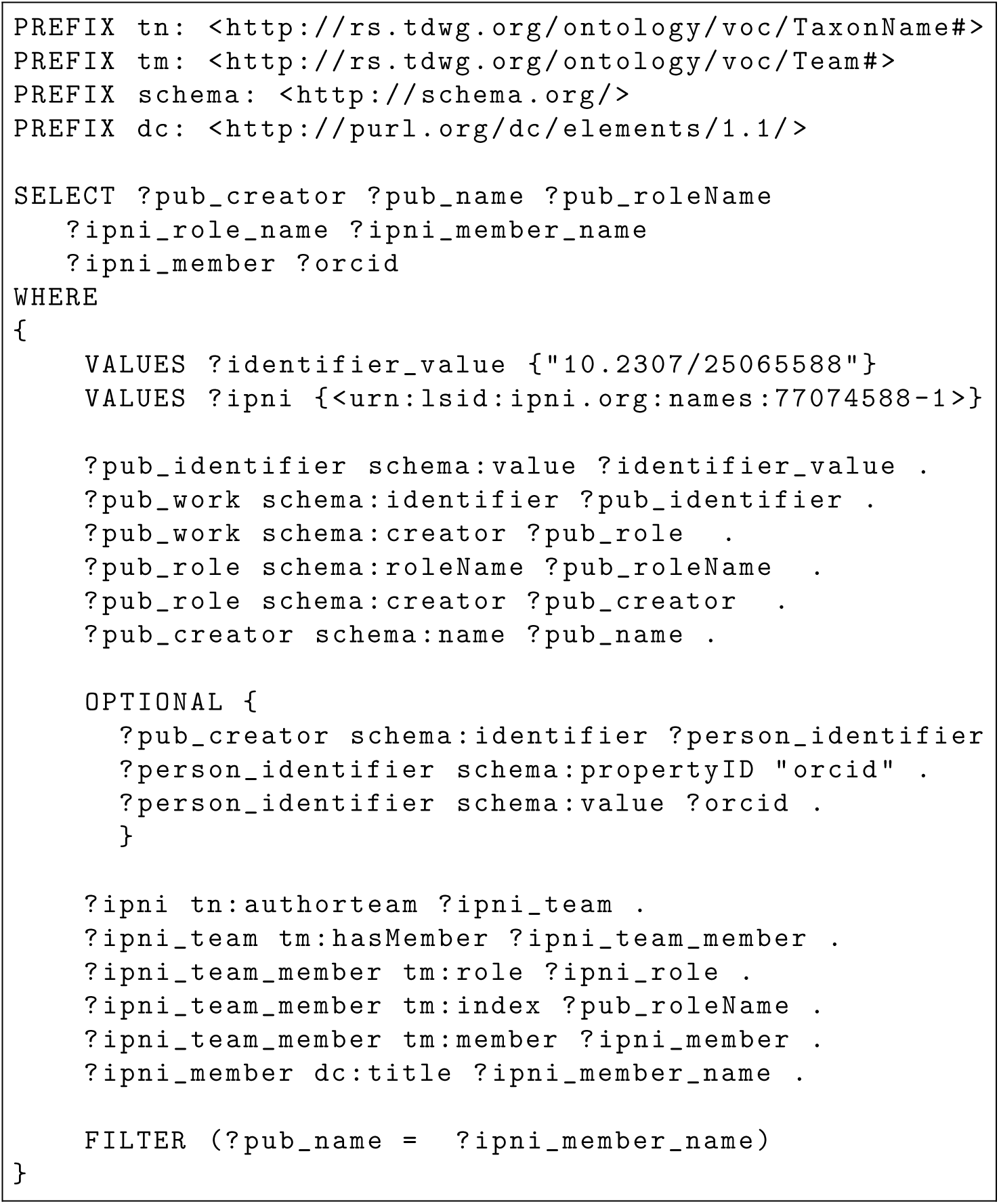
SPARQL query to match author in IPNI and ORCID

## Footnotes

1 http://www.ipni.org

2 https://blazegraph.com

3 https://biodiversitylibrary.org

4 https://github.com/rdmpage/ipni-names

5 https://crossref.org

6 https://orcid.org

7 http://schema.org

8 https://www.wikidata.org

9 https://www.ncbi.nlm.nih.gov/genbank/

10 https://treebase.org

11 https://gbif.org

## References

1. Bik, H.M.: Let’s rise up to unite taxonomy and technology. PLOS Biology 15(8), e2002231 (Aug 2017). https://doi.org/10.1371/journal.pbio.2002231

2. Clark, T., Martin, S., Liefeld, T.: Globally distributed object identification for biological knowledgebases. Briefings in Bioinformatics 5(1), 59–70 (Mar 2004). https://doi.org/10.1093/bib/5.1.59

3. Croft, J., Cross, N., Hinchcliffe, S., Lughadha, E.N., Stevens, P.F., West, J.G., Whitbread, G.: Plant names for the 21st century: the International Plant Names Index, a distributed data source of general accessibility. TAXON 48(2), 317–324 (1999). https://doi.org/10.2307/1224436

4. Franz, N.M., Sterner, B.W.: To increase trust, change the social design behind aggregated biodiversity data. Database 2018 (Jan 2018). https://doi.org/10.1093/database/bax100

5. Holland, V.T., Johnson, J.: Introducing ’Role’ (2014), http://blog.schema.org/2014/06/introducing-role.html

6. King, D., Morse, D., Willis, A., Dil, A.: Towards the bibliography of life. ZooKeys 150, 151–166 (Nov 2011). https://doi.org/10.3897/zookeys.150.2167, https://zookeys.pensoft.net/article/3035/

7. Konno, T.U.P., Rapini, A., Goyder, D.J., Chase, M.W.: The new genus Minaria (Asclepiadoideae, Apocynaceae). TAXON 55(2), 421–430 (2006). https://doi.org/10.2307/25065588

8. Page, R.D.M.: LSID Tester, a tool for testing Life Science Identifier resolution services. Source Code for Biology and Medicine 3(1), 2 (Feb 2008). https://doi.org/10.1186/1751-0473-3-2

9. Page, R.D.M.: Towards a biodiversity knowledge graph. Research Ideas and Outcomes 2, e8767 (Apr 2016). https://doi.org/10.3897/rio.2.e8767

10. Page, R.D.M.: Liberating links between datasets using lightweight data publishing: an example using plant names and the taxonomic literature. Biodiversity Data Journal (6) (Jul 2018). https://doi.org/10.3897/BDJ.6.e27539

11. Page, R.D.M.: Ozymandias: a biodiversity knowledge graph. PeerJ 7, e6739 (Apr 2019). https://doi.org/10.7717/peerj.6739

12. Penev, L., Dimitrova, M., Senderov, V., Zhelezov, G., Georgiev, T., Stoev, P., Simov, K.: OpenBiodiv: A Knowledge Graph for Literature-Extracted Linked Open Data in Biodiversity Science. Publications 7(2), 38 (May 2019). https://doi.org/10.3390/publications7020038

13. Penev, L., Paton, A., Nicolson, N., Kirk, P., Pyle, R.L., Whitton, R., Georgiev, T., Barker, C., Hopkins, C., Robert, V., Biserkov, J., Stoev, P.: A common registration-to-publication automated pipeline for nomenclatural acts for higher plants (International Plant Names Index, IPNI), fungi (Index Fungorum, MycoBank) and animals (ZooBank). ZooKeys (550), 233–246 (Jan 2016). https://doi.org/10.3897/zookeys.550.9551

14. Shorthouse, D., Page, R.D.M.: Quantifying Institutional Reach Through the Human Network in Natural History Collections. Biodiversity Information Science and Standards 3, e35243 (Aug 2019). https://doi.org/10.3897/biss.3.35243

15. Vrandečcić, D., Krötzsch, M.: Wikidata. Communications of the ACM 57(10), 78–85 (Sep 2014). https://doi.org/10.1145/2629489

